# Profiling cell proliferation after whole-genome duplication in human cells

**DOI:** 10.64898/2026.03.12.711482

**Authors:** Guang Yang, Masaya Inoko, Kaito Ogura, Sumire Ishida-Ishihara, Yuki Tsukada, Akira Funahashi, Masanao Sato, Ryota Uehara

## Abstract

Though whole-genome duplication (WGD) contributes to cancer progression, the mechanism of post-WGD cell proliferation remains unclear. Here, using 6-day live-imaging, we analyzed the proliferation dynamics of more than 150 post-WGD HCT116 cell lineages. A quantitative comparison of mitotic patterns and cell fates between proliferative and non-proliferative lineages revealed that multipolar chromosome segregation in early mitosis is a key factor limiting the proliferative capacity of post-WGD progenies. Multipolar chromosome segregation suppressed post-WGD cell viability, particularly when accompanied by drastic chromosome loss or when it repeatedly occurred. Tracing proliferative lineages elucidated that they proliferated mainly by imposing the risk of multipolar chromosome segregation on one of two sub-lineages that formed after the first bipolar division. Meanwhile, a considerable proportion of proliferative lineages consisted entirely of progeny of early multipolar chromosome segregation events. Our results highlight key cellular events that determine the proliferation dynamics and diversity of post-WGD progenies, providing a fundamental reference for understanding WGD-associated bioprocesses.

**Summary statement:** Live image tracing of >150 cell lineages reveals the cross-generation dynamics of multipolar chromosome segregation that determine the fates of post-whole-genome duplication progeny cells.

## Introduction

Whole-genome duplication (WGD) occurs when a cell skips chromosome segregation after DNA replication, resulting in the doubling of the entire cell component. More than 30% of solid tumors commonly experience at least one round of WGD (Bielski et al., 2018), potentially contributing to cancer progression in broad tissue backgrounds (Fujiwara et al., 2005; Vittoria et al., 2023). Therefore, understanding the key determinants of post-WGD cell survival and proliferation has become an emerging important topic in biological and medical research (Quinton et al., 2021; Vittoria et al., 2023; Yoshizawa et al., 2023). However, WGD leads to chromosome instability (CIN) and highly heterogeneous genome abnormalities in their progenies (Kuznetsova et al., 2015; Prasad et al., 2022). This process is thought to confer various properties and fates on post-WGD cells, making it difficult to develop universal approaches to selectively suppress them.

CIN in post-WGD cells has been largely attributed to mitotic defects caused by the doubled number (i.e., 4) of centrosomes resulting from WGD (Godinho et al., 2009). Possession of more than 2 centrosomes increases the risk of multipolar spindle formation, leading to unequal chromosome segregation and endangering genome integrity in progeny cells. To manage the risk posed by supernumerary centrosome formation, cells have mechanisms to cluster excess centrosomes into two spindle poles (Basto et al., 2008; Ganem et al., 2009; Kwon et al., 2008; Silkworth et al., 2009). The centrosome clustering mechanism operates in post-WGD cells and is thought to contribute significantly to their survival (Ganem et al., 2009). Moreover, a recent study has revealed that the 4 centrosomes in post-WGD cells are sometimes segregated into two spindle poles in a 1:3 ratio, contributing to the rapid removal of excess centrosomes in one of their daughter cells (Baudoin et al., 2020). However, the above spindle bipolarization occurs only in a limited fraction of post-WGD cells, in a cell-type-dependent manner (Lau et al., 2024). Therefore, a substantial proportion of post-WGD cells suffer multipolar chromosome segregation, often leading to cell death in the subsequent cell cycle (Ganem et al., 2009; Inoko et al., 2026). As a result, very few post-WGD cells acquire survival and enter metastable proliferation in the long term (Inoko et al., 2026). Previous studies have analyzed post-WGD cell proliferation either by tracing a few cell generations after WGD (11, 13) or by end-point population assays such as colony formation (Inoko et al., 2026; Vora et al., 2025). For this reason, the long-term dynamics of the formation of surviving post-WGD cell lineages, including potential generation-wise changes in cell cycle progression or mitotic fidelity, have been largely unknown. This limitation of knowledge has precluded the understanding of key determinants of post-WGD cell viability.

In this study, we investigated cell proliferation dynamics across several generations after WGD in human colorectal near-diploid HCT116 cells, quantitatively addressing the characteristics of various cell lineages to understand the potential determinants of the long-term fates of post-WGD cells.

## Results and discussion

### Variation in proliferative ability among different post-WGD cell lineages

To gain insight into the post-WGD cell proliferation principle, we induced WGD in HCT116 cells via cytokinesis failure using cytochalasin B (Fig. 1A; see Materials and methods for details). The subsequent proliferation of WGD-induced cells was live-imaged for 6 d. Interphase nuclei and mitotic chromosomes were visualized by stable expression of histone H2B-mCherry (Fig. 1B). WGD-induced cells were readily distinguished from uninduced cells based on possession of 2 nuclei during the first cell cycle after cytochalasin B treatment (Fig. 1B). In entire view of cell culture well after 6 d live-imaging, we found colonies with various sizes originated from a limited proportion of WGD cells (Fig. 1B; Movie S1-3). This finding indicates highly heterogeneous fates of post-WGD lineages, with a very small chance of sustainable proliferation.

**Fig. 1:**
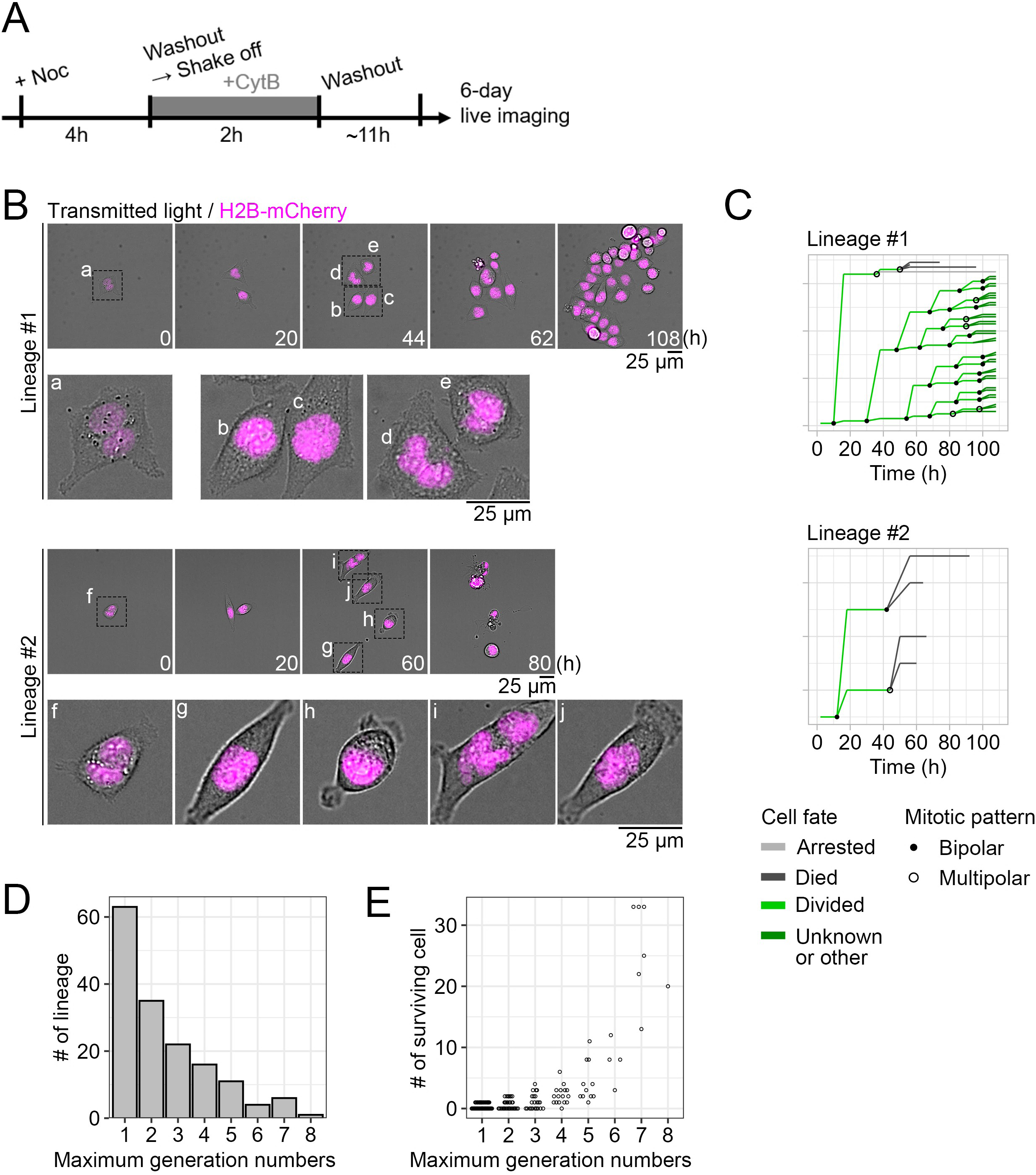
Heterogeneity in proliferative capacity among post-WGD cell lineages. **(A)** Scheme of WGD induction and subsequent live-imaging. **(B)** Examples of live images of post-WGD cell proliferation. Images were taken at a 2-h interval. The entire view of the lineage and magnified views of individual cells are shown in the top and bottom, respectively, for each example lineage. The initial post-WGD cells are shown as a and f. Representative cells that experienced bipolar (b, c, g, or h) or multipolar chromosome segregation (d, e, i, or j) in the previous mitosis are also shown. **(C)** Examples of post-WGD cell lineage tracings in B. Makers indicated different mitotic patterns. Progenies were color-coded by cell fate. The tracings of all 158 post-WGD cell lineages are provided as Supplementary material S1. **(D)** Maximum generation numbers that individual post-WGD cell lineages reached within 108 h of live-imaging in B. 158 lineages from 3 independent experiments. **(E)** Distribution of surviving cell numbers within individual lineages, plotted against their maximum cell generations, at 108 h after the initiation of live-imaging in B.

To analyze the above trend quantitatively, we manually tracked 158 post-WGD cell lineages in an unbiased manner, annotated fates of their progenies (categorized as “died,” “arrested,” or “divided”) and patterns of their mitotic events (“bipolar” or “multipolar”), and obtained their proliferation profiles (Fig. 1C; see Materials and methods for details). From these profiles, we quantified maximum cell generations that individual lineages reached (Fig. 1D) and numbers of their survived progenies at the end of the analyzed time window (5 d; Fig. 1E). While 40% (63 out of 158) of post-WGD cell lineages did not reach the second cell generation, a small proportion of them (8%; 11 out of 158) reached more than fifth generation with the maximum generation of 8 (Fig. 1D). Lineages reaching more generations tended to contain more survived progenies (Fig. 1E). However, even in the most proliferative lineages, numbers of survived progenies were much less than those expected from the exponential cell number increase (Fig. 1E). These quantifications collectively demonstrate defective and heterogenous nature of post-WGD cell proliferation in HCT116 cells.

### Early cell death limits the proliferative capacity of post-WGD cell lineages

We wished to gain further insight into the determinants of proliferative capacity of post-WGD cell lineages. For this, we categorized the post-WGD cell lineages analyzed above based on their survived progenies numbers: We defined the lineages with ≥15 survived progenies as “proliferative”, and those with <15 survived progenies as “non-proliferative”, and compared detailed profiles between these groups (Fig. 2A). We found a prominent difference in the distribution of early progeny fates between these groups: While 17.8% or 35.9% of the first- or second-generation progenies suffered cell death, respectively, in non-proliferative lineage group, all progenies survived through the first and second generation in proliferative lineage group (Fig. 2A). While viability (proportion of “divided” progenies) decreased as generation increase in proliferative lineage group, more progenies still survived in this group compared to non-proliferative lineage group even after the second generation (Fig. 2A). We also analyzed cell cycle length, quantified from progenies surviving through the corresponding cell generation, in these lineage groups (Fig. 2B; note that cell cycle length in the first generation was measured as time taken from the initiation of cytochalasin B treatment to the entry into the first mitosis). Interestingly, in both proliferative and non-proliferative lineage groups, cell cycle length after the second generation became drastically shorter than that in the first generation (Fig. 2B). This data suggests that post-WGD cells overcome WGD-associated cell cycle delay after the first mitosis, possibly through adaptive mechanisms. Meanwhile, cell cycle length tended to be shorter in the proliferative lineage group than in the non-proliferative lineage group, though their difference was not statistically significant in most cell generations (Fig. 2B). Overall, more proliferative post-WGD cell lineages show prominently higher cell survival, particularly in the earlier phase after WGD, compared to those with less proliferative capacity.

**Fig. 2:**
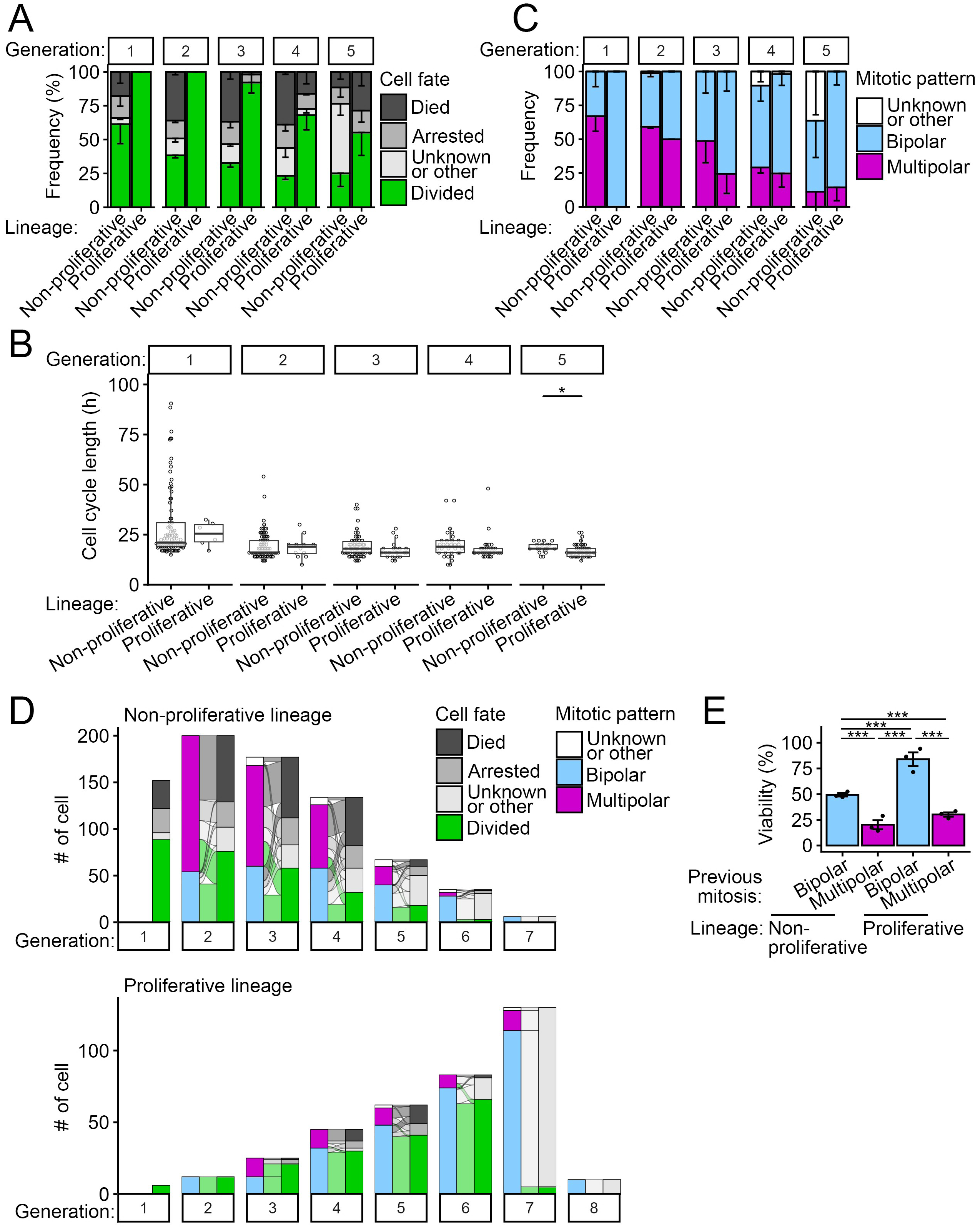
Multipolar chromosome segregation limits the viability of post-WGD progenies. **(A)** Frequency of cell fates of post-WGD progenies in proliferative or non-proliferative lineages, sorted by cell generation. Mean ± SE of 3 independent experiments. In total, 150 or 730 cells from 6 proliferative or 152 non-proliferative lineages, respectively, from 3 independent experiments were analyzed. **(B)** Cell cycle length, measured for viable cells, sorted by generations, of proliferative or non-proliferative lineages. Note that the cell cycle length in the first generation was quantified as the duration from the initiation of cytochalasin B treatment to the entry into the first mitosis. In total, 110 or 273 cells from 6 proliferative or 89 non-proliferative lineages, respectively, from three independent experiments were analyzed. Asterisks indicate a statistically significant difference between the lineage groups (*p < 0.05, Brunner-Munzel test). **(C)** Frequency of mitotic pattern in proliferative or non-proliferative lineages in each round of mitosis. Mean ± SE of 3 independent experiments. In total, 110 or 273 cells from 6 proliferative or 89 non-proliferative lineages, respectively, from 3 independent experiments were analyzed. **(D)** Alluvial diagrams showing the relationship between chromosome segregation pattern in each mitotic event and the fate of the daughter cell in the subsequent cell cycle. Profiles of mitotic patterns and cell fates were pooled from 6 proliferative or 152 non-proliferative lineages. **(E)** Viability of post-WGD progenies immediately after bipolar or multipolar chromosome segregation. Mean ± SE of 3 independent experiments. Asterisks indicate statistically significant differences among samples (***p < 0.001, the Steel-Dwass test).

### Multipolar chromosome segregation as a primary cause of early cell death in post-WGD lineage

We next compared the dynamics of mitotic patterns across generations between the proliferative and non-proliferative lineage groups (Fig. 2C). Their mitotic patterns were most distinct in the first mitosis: While 67% of non-proliferative lineages underwent multipolar chromosome segregation, all proliferative lineages underwent bipolar chromosome segregation in the first mitosis (Fig. 2C). In the second mitosis, frequency of multipolar chromosome segregation drastically increased in proliferative lineages and became equivalent to that in non-proliferative lineages. This result indicates that bipolarization of the chromosome segregation pattern at the first mitosis is a key determinant of the survivability of post-WGD cell lineages. After the third generation, the proportion of multipolar chromosome segregation gradually decreased in both proliferative and non-proliferative lineages, suggesting a gradual reduction in the risk of multipolar chromosome segregation. This trend is consistent with the gradual removal of excess centrosomes from the post-WGD cell population in previous studies (Baudoin et al., 2020; Ganem et al., 2009; Godinho et al., 2014; Kuznetsova et al., 2015; Potapova et al., 2016; Yaguchi et al., 2018).

To further address the impacts of mitotic patterns on proliferative quality of post-WGD cell lineages, we directly correlated the mitotic patterns and subsequent daughter cell fates in each generation, either in proliferative or non-proliferative lineages (Fig. 2D). Alluvial diagrams in each generation clearly show that, across generations, daughter cell death or arrest mainly occurred following the previous multipolar chromosome segregation. Overall, 65.8% or 73.7% of non-proliferative daughters (i.e., died or arrested) experienced multipolar chromosome segregation in the immediate past mitotic events across 2-5 generations in proliferative or non-proliferative lineages, respectively. This data indicates that multipolar chromosome segregation is the primary cause of non-proliferative cell fate in post-WGD cell lineages.

On the other hand, the data also show that a substantial proportion of daughter cells survived through multipolar chromosome segregation. For example, 85.2% or 24.9% of the third-generation progenies formed through multipolar chromosome segregation in the second mitosis survived to undergo the next mitosis in proliferative or non-proliferative lineages, respectively (Fig. 2D). Quantification of cell viability across 1-5 generations also demonstrates overall 30.1% or 20.2% of multipolar chromosome segregation events resulted in viable daughters in proliferative or non-proliferative lineages, while bipolar chromosome segregation resulted in significantly more frequent viable daughters (Fig. 2E). Therefore, the frequent multipolar chromosome segregation largely explains the limited proliferative ability of the post-WGD cell lineages. Meanwhile, our data also suggest that multipolar chromosome segregation can contribute to the generation of surviving post-WGD progenies, potentially fueling the genome complexity of post-WGD sub-lineages.

### Chromosome loss upon multipolar chromosome segregation limits post-WGD cell viability

Since multipolar chromosome segregation was the key factor limiting the proliferative capacity of post-WGD progenies, we further addressed aspects of daughter cells affected by these events. For this, we compared the patterns of chromosome distribution upon bipolar or multipolar chromosome segregation by quantifying the ratio of nuclear H2B-mCherry signals among daughter cells immediately after individual mitotic events (Fig. 3A). Consistent with previous studies (Gisselsson et al., 2010; Inoko et al., 2026), daughters cells generated through multipolar chromosome segregation (including tripolar and tetrapolar segregation) often got fused with one another, resulting in multinuclear daughter cells (Fig. 3A). Therefore, we separately analyzed chromosome segregation ratio for mitotic events generating different numbers of daughters (Fig. 3B). Bipolar chromosome segregation resulted in nearly equal chromosome distribution between two daughters, resulting in on average 0.5 distribution ratio with a very small deviation (Fig. 3B). Multipolar chromosome segregation generating more than 2 daughters naturally resulted in reduction in the average chromosome distribution ratio with a drastically high deviation compared to bipolar control (Fig. 3B). Even when multipolar chromosome segregation generated 2 daughters, chromosome distribution between them was highly deviated, often resulting in prominent gain and loss of chromosome content (Fig. 3B). This result demonstrates that multipolar chromosome segregation is a primary source of drastic changes in chromosome contents in post-WGD cell populations.

**Fig. 3:**
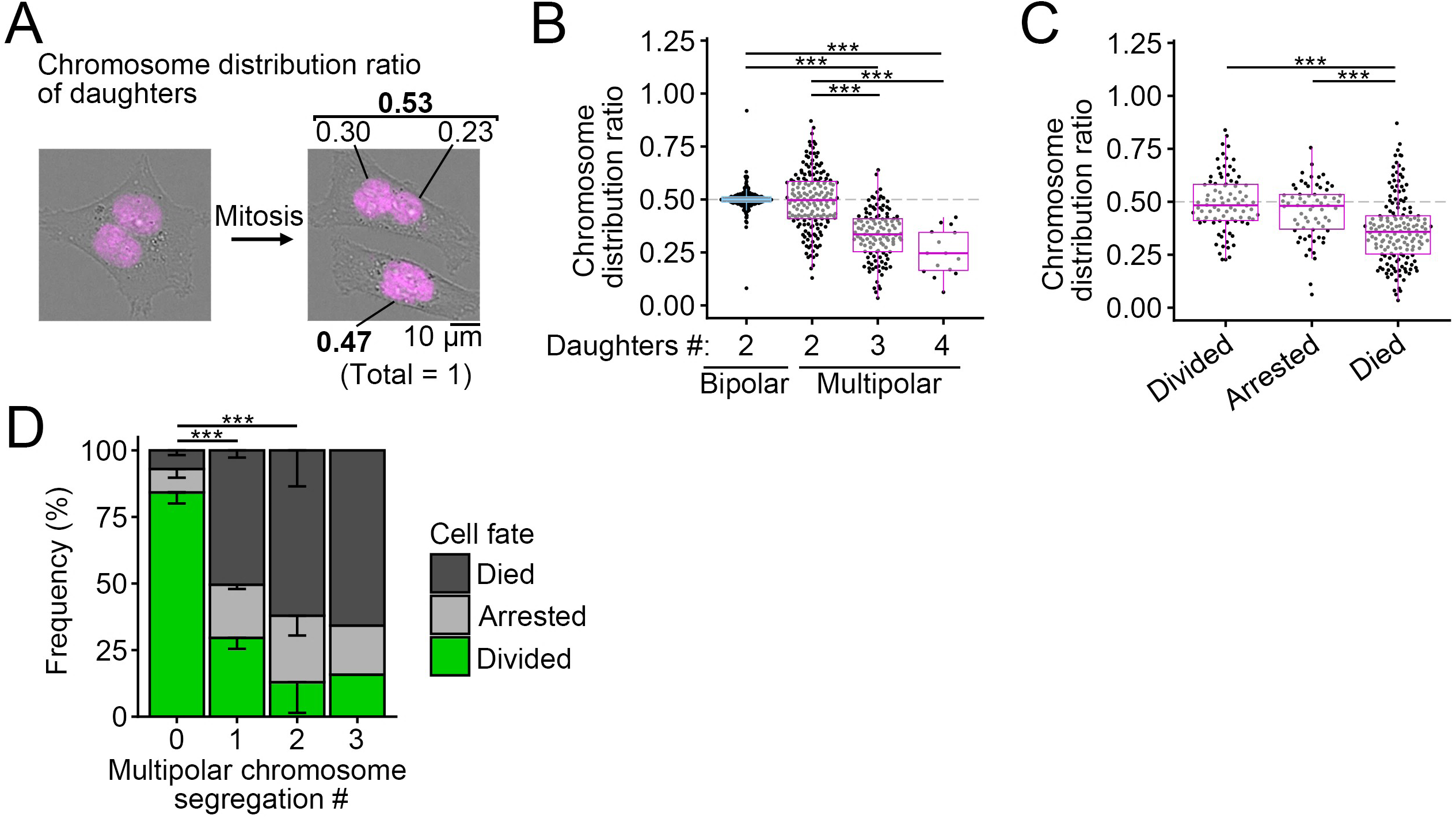
Drastic changes in chromosome contents upon multipolar chromosome segregation. **(A)** Scheme of quantification of chromosome distribution ratio based on segregation ratio of nuclear H2B-mCherry intensity among daughter cells. **(B, C)** Chromosome distribution ratio in each daughter cell, sorted by previous mitotic pattern and daughter cell number (B), or daughter cell fate (C). At least 15 or 66 cells from 3 independent experiments for each condition in B or C, respectively. Asterisks indicate statistically significant differences among samples (***p < 0.001, the Steel-Dwass test). **(D)** Frequency of fates of post-WGD progenies, sorted by the number of repetitions of multipolar chromosome segregation. Mean ± SE of 3 independent experiments (at least 81 cells were analyzed for each condition), except for the condition with 3 repetitions of multipolar chromosome segregation (20 cells from 2 independent experiments). Asterisks indicate statistically significant differences in the frequency of divided cells among samples (***p < 0.001, the Steel-Dwass test).

We next analyzed the relationship between chromosome distribution ratio in daughter cells forming through multipolar chromosome segregation and fates of these daughters (Fig. 3C). Daughter cells that survived through their generations showed average chromosome distribution ratio of 0.49 upon the immediate past mitosis (Fig. 3C), which was close to the ratio observed for bipolar chromosome segregation (Fig. 3B). Importantly, inviable daughters (categorized as “died”) showed significantly lower chromosome distribution ratio than viable daughters (Fig. 3C), indicating that a drastic chromosome loss caused by unequal chromosome segregation upon multipolar division promotes subsequent cell death to limit proliferative capacity of post-WGD cell lineages.

### Cumulative effects of multipolar chromosome segregation on post-WGD cell viability

While multipolar chromosome segregation often led to subsequent daughter cell death, a substantial proportion of daughters remained viable even after multipolar chromosome segregation (Fig. 2D). This led to sub-lineages undergoing multiple rounds of multipolar chromosome segregation over generations. To quantitatively understand the effects of repetitive multipolar chromosome segregation on progenies’ fates, we sorted the viability of post-WGD progenies based on the number of multipolar chromosome segregations they had experienced (Fig. 3D). We observed that the post-WGD progenies experienced at most 3 repetitions of multipolar chromosome segregation in 2 out of 3 independent experiments. While 29.6% of progenies remained viable after one round of multipolar chromosome segregation, viability gradually decreased as the number of repetitions increased. This result demonstrates the cumulative adverse effects of multipolar chromosome segregation on post-WGD cell proliferation. However, the data also show that a considerable proportion of progenies remained viable after multiple rounds of multipolar chromosome segregation (Fig. 3D). Considering that multipolar chromosome segregation induces drastic changes in chromosome contents (Fig. 3B), multiple rounds of multipolar chromosome segregation might be an important principle that causes drastic structural and copy number abnormalities in post-WGD cell lineages (Prasad et al., 2022).

### Diversity in mitotic patterns among proliferative post-WGD cell lineages

Finally, we wished to understand how post-WGD cell lineages manage or accept the adverse effects of multipolar chromosome segregation to achieve proliferation. For this, we selected 12 post-WGD cell lineages from the entire view of culture wells that formed relatively large colonies at the end of live imaging and analyzed their mitotic patterns throughout colony formation (until 108 or 140 h after the initiation of live imaging; Fig. 4A-L). We found two major patterns of proliferative lineage formation. i) In the most prevalent case (7 out of 12 lineages; categorized as “asymmetric lineage”; Fig. 4A-G), bipolar chromosome segregation occurred in the first mitosis, followed by asymmetric fates of the resulting two sub-lineages: While one sub-lineage underwent nearly lethal multipolar chromosome segregation in the second mitosis, the other sub-lineage readily proliferated by repeating bipolar chromosome segregation in most of subsequent mitotic events. This mitotic pattern resulted in a situation in which only one of two sub-lineages, formed in the first mitosis, crucially contributed to the formation of post-WGD progenies. ii) In the next most common case (3 out of 12 lineages; “ex-multipolar lineage”; Fig. 4H-J), all surviving progenies formed through at least one round of multipolar chromosome segregation. iii) Of the remaining 2 lineages, one showed relatively symmetric growth of two sub-lineages formed by bipolar chromosome segregation at the first mitosis, followed by occasional multipolar chromosome segregation in later mitotic events (Fig. 4K). The last lineage also underwent bipolar chromosome segregation at the first mitosis. However, one of its sub-lineages became lethal in the third generation, resulting in an asymmetric growth pattern of the sub-lineages (Fig. 4L).

**Fig. 4:**
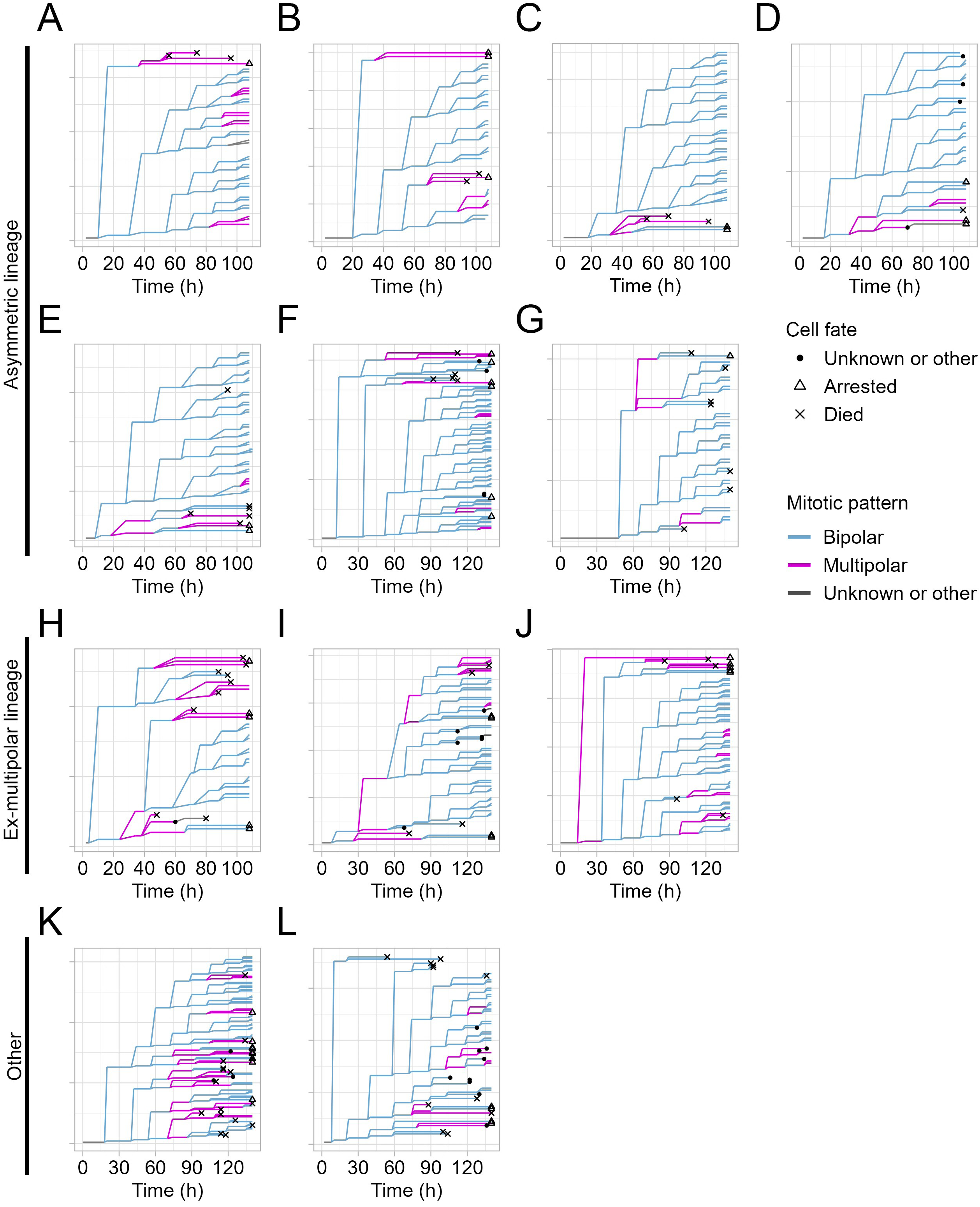
Diversity in mitotic patterns among proliferative post-WGD cell lineages. **(A-L)** Tracings of proliferative post-WGD cell lineages throughout the live-imaging. Progenies were color-coded according to mitotic patterns in the previous mitosis. Makers indicated the fates of progenies. The tracing in A is identical to that in Fig. 1C. Note that lineages in F and I-L were newly selected from the entire view of cell culture wells, and not used in the above analyses in Fig. 1-3.

The above lineage analysis revealed prominent diversity in the proliferation dynamics of post-WGD progenies, which could be classified into a few distinct patterns. The highest prevalence of “asymmetric lineages” indicates that this mitotic pattern is the most effective strategy for post-WGD cell proliferation, at least in the HCT116 background (Fig. 4A-G). In this case, the risk of multipolar chromosome segregation is imposed on one of the sub-lineages arising from the first bipolar division. A possible mechanism to achieve these proliferation patterns would be the asymmetric distribution of excess centrosomes formed upon WGD, as reported in a previous study (13): Once one side of a post-WGD cell lineage loses excess centrosomes, it becomes able to undergo stable bipolar divisions in subsequent generations. On the other hand, a considerable proportion of lineages consisted entirely of progenies formed through multipolar chromosome segregation (Fig. 4H-J), demonstrating the unignorable contribution of multipolar chromosome segregation to the characterization of post-WGD cell lineages.

In this study, we attempted to identify determinants of post-WGD cell fates through comprehensive lineage tracking. We found that alterations in chromosome content due to multipolar chromosome segregation and the magnitudes of its repetition are key factors limiting the viability of post-WGD progenies across generations. While post-WGD lineages acquired proliferative capacity by ensuring long-term mitotic stability by avoiding the risk of multipolar chromosome segregation in the earlier phase of their formation, a substantial proportion of lineages also form under the influence of erroneous multipolar divisions. To achieve rational control of post-WGD cell proliferation, it will be crucial in future studies to elucidate the correspondence between the above-described mitotic patterns and the resulting genomic compositions or gene expression profiles in individual post-WGD subclones. Our results provide a fundamental reference for further discussion of the principles governing the diversification of WGD cell populations across different bioprocesses.

## Materials and methods

### Cell culture

HCT116 cells (WT line, RRID: CVCL_0291 from Riken BRC) were cultured in McCoy’s 5A (Wako) supplemented with 10% fetal bovine serum (FBS) and 1× antibiotic-antimycotic solution (AA; Sigma-Aldrich) at 37°C with 5% CO2. Transgenic cell lines were established by transfecting HCT116 cells with plasmids encoding histone H2B-mCherry (Inoko et al., 2026) using JETPEI (Polyplus-transfection), followed by selection of positive clones with the appropriate antibiotics.

To induce WGD, cells were first arrested at prometaphase by treating with 40 ng/mL nocodazole for 4 h, washing 3 times with supplemented culture medium, and then shaking off. These shaken-off mitotic cells were treated with 5 μg/mL cytochalasin B for 2 h, followed by washing 3 times with supplemented culture medium.

### Live cell imaging

For live imaging of cell proliferation after WGD induction, we seeded WGD-induced cells on an 8-well cover glass-bottom chamber (zell-kontakt GmbH), replaced culture media with supplemented phenol red-free McCoy’s 5A (Cytiva) at 4 h after WGD induction, and started cell imaging from 12.5 or 13 h after WGD induction. Live imaging was conducted at 37°C with 5% CO2 using a Ti2 microscope with a ×20 0.75 NA Plan-Apochromatic objective (Nikon) and a Zyla4.2 sCMOS camera (Andor). Image acquisition was controlled by µManager.

For unbiased cell lineage tracking, we analyzed all post-WGD cell lineages in approximately one-fifth of the entire well in each of three independent experimental trials. For lineage tracing in Fig. 4, we also analyzed 5 lineages outside the well areas described above. Cell fates and mitotic events in all cell lineages were manually analyzed using ImageJ (NIH).

### Statistic analysis

Analyses for significant differences between the two groups were conducted using the Brunner-Munzel test in R software (The R Foundation). Multiple group analyses were conducted using the Steel-Dwass test in R software. Statistical significance was set at p < 0.05. P-values are indicated in figures or the corresponding figure legends.

## Author Contributions

Conceptualization, G.Y., and R.U.; Methodology, G.Y., M.I., Y.T., A.F., M.S., and R.U.; Investigation, G.Y., and M.I.; Formal Analysis, G.Y., M.I., K.O., and R.U.; Resources, G.Y., M.I., S.I., Y.T., A.F., M.S., and R.U.; Supervision, R.U.; Writing – Original Draft, R.U.; Writing – Review & Editing, G.Y., and R.U.; Funding Acquisition, G.Y., S.I., and R.U.

## Data and resource availability

All relevant data and details of resources can be found within the article and its supplementary information.

---

**Supplementary movie S1-3: Entire views of all post-WGD cell live images**

Live images of post-WGD cell proliferation (2 h per frame). Data from 3 independent experiments are shown in separate movie files. The stitched field of view is 1 cm × 1 cm.

**Supplementary material S1: Tracing of all post-WGD cell lineages**

Makers indicated different mitotic patterns. Progenies were color-coded based on their fates.

