## Supplementary Material S1 for "Profiling cell proliferation after whole-genome duplication in human cells"

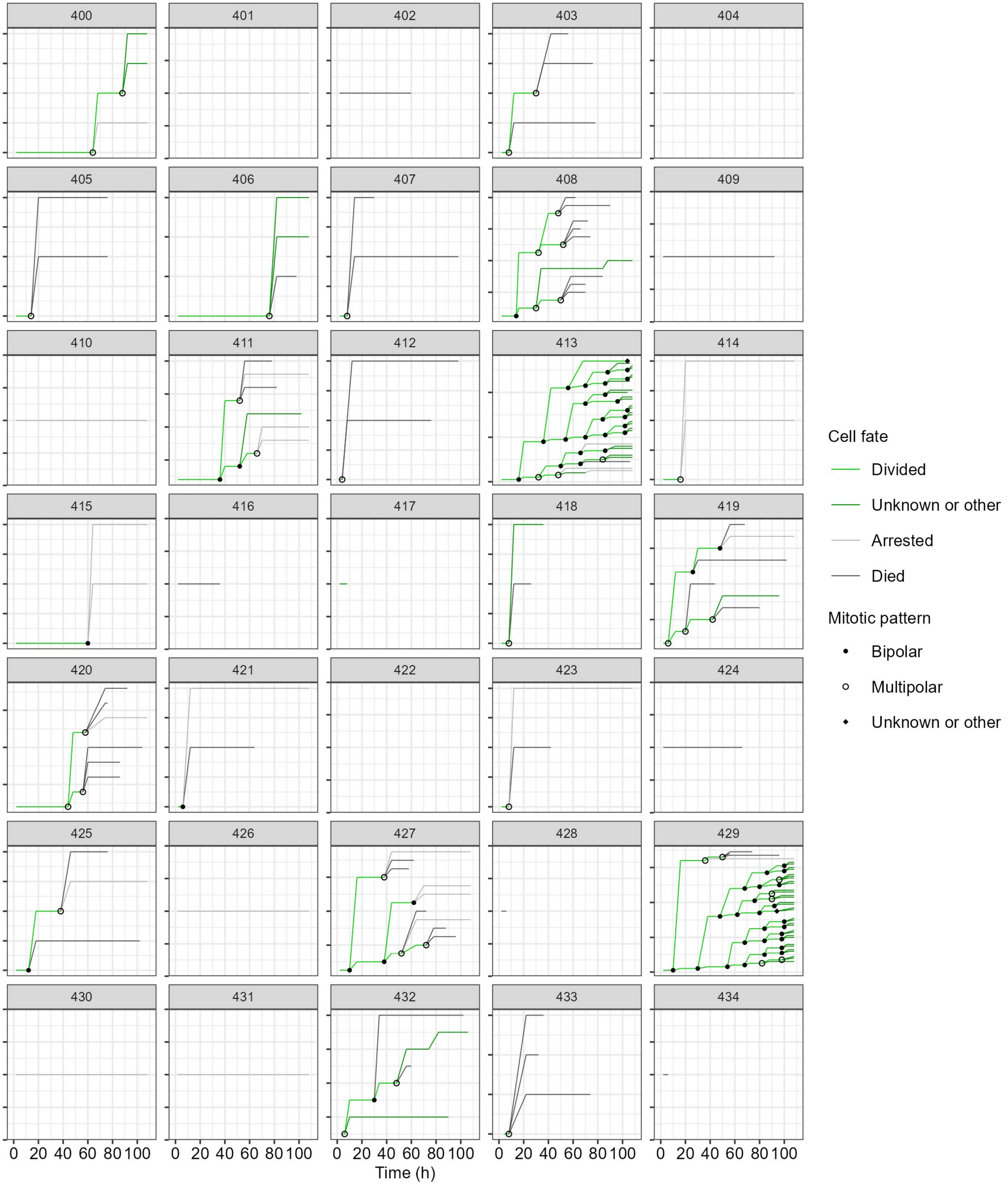

Supplementary material S1 (continued)

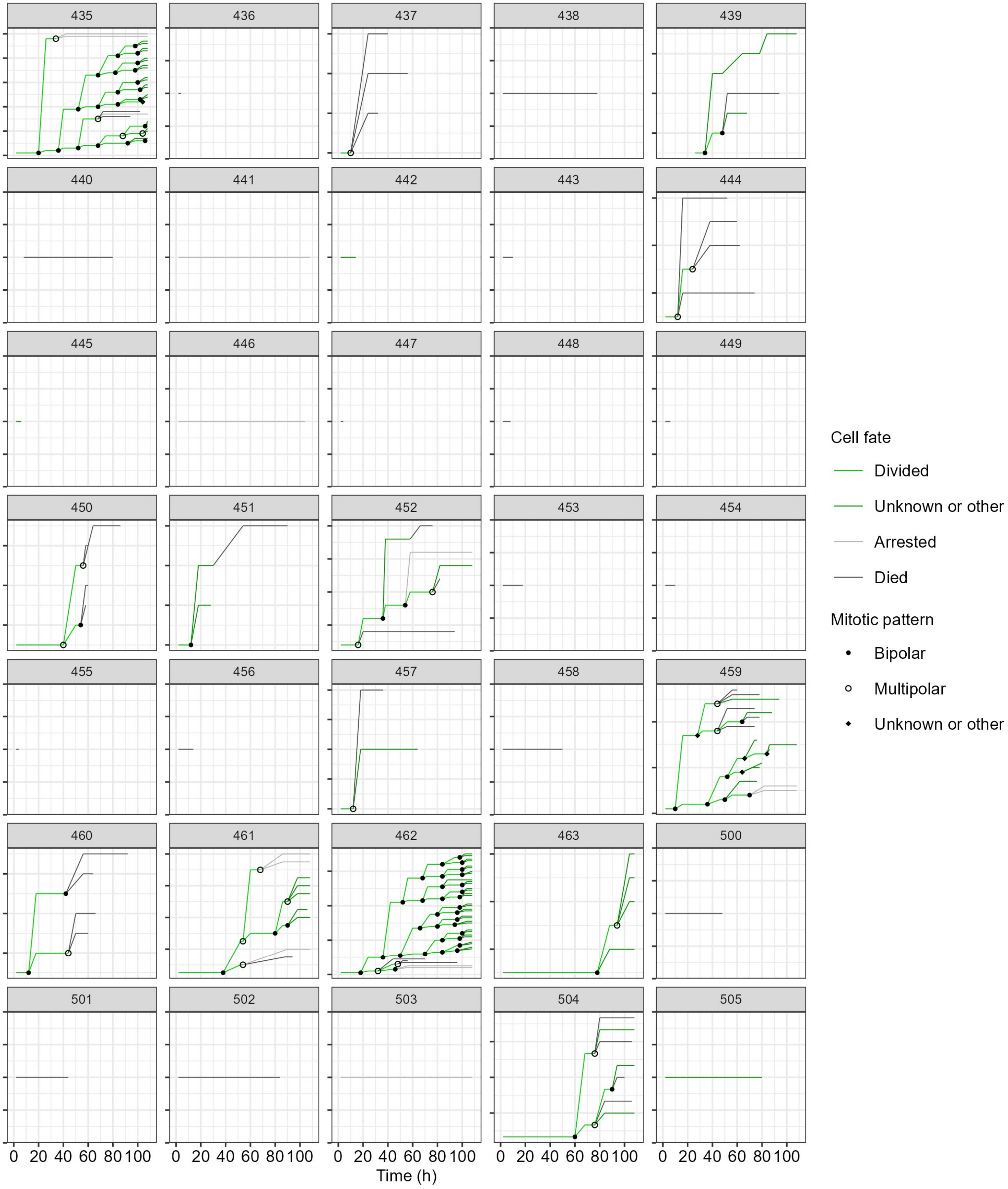

Supplementary material S1 (continued)

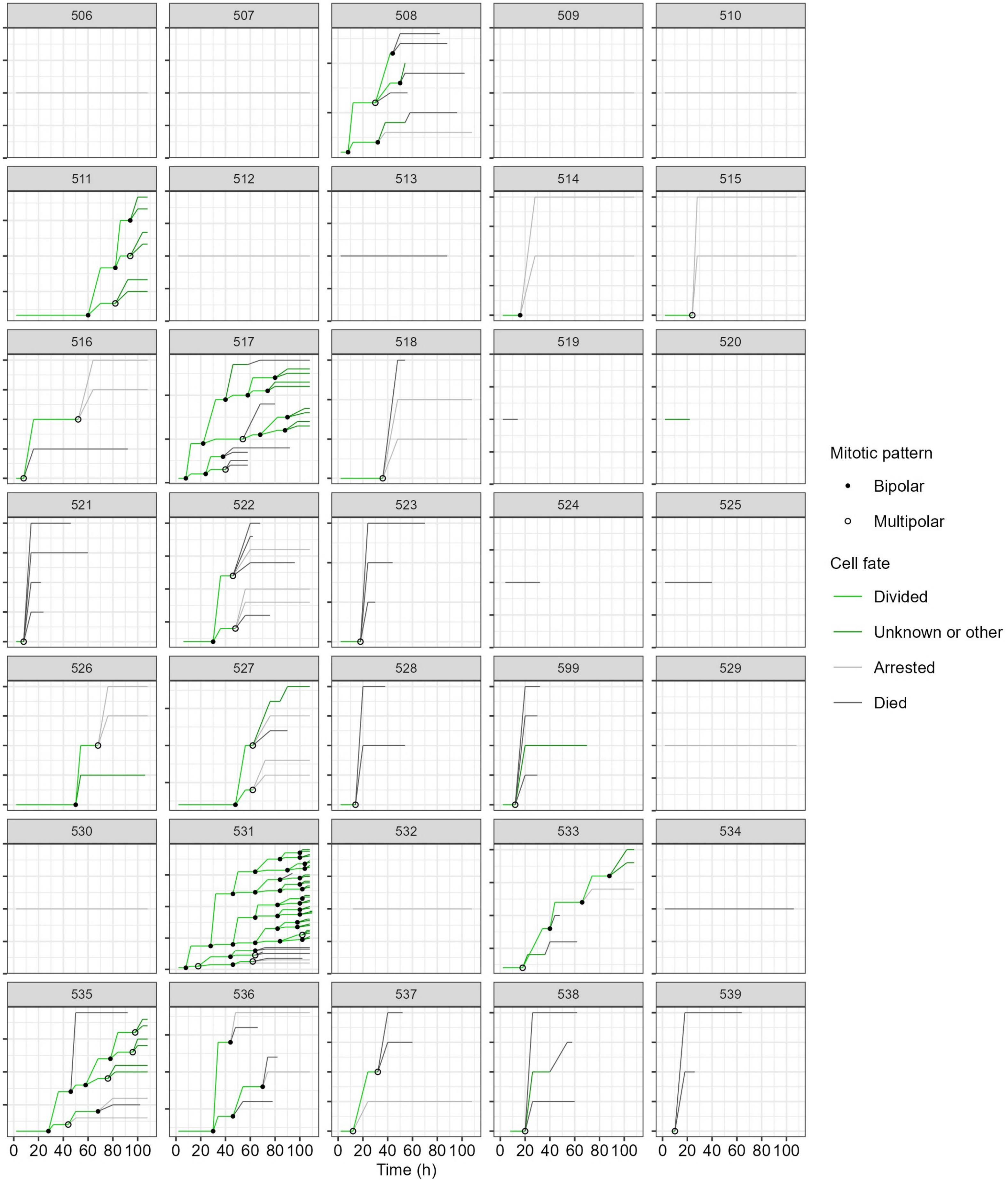

Supplementary material S1 (continued)

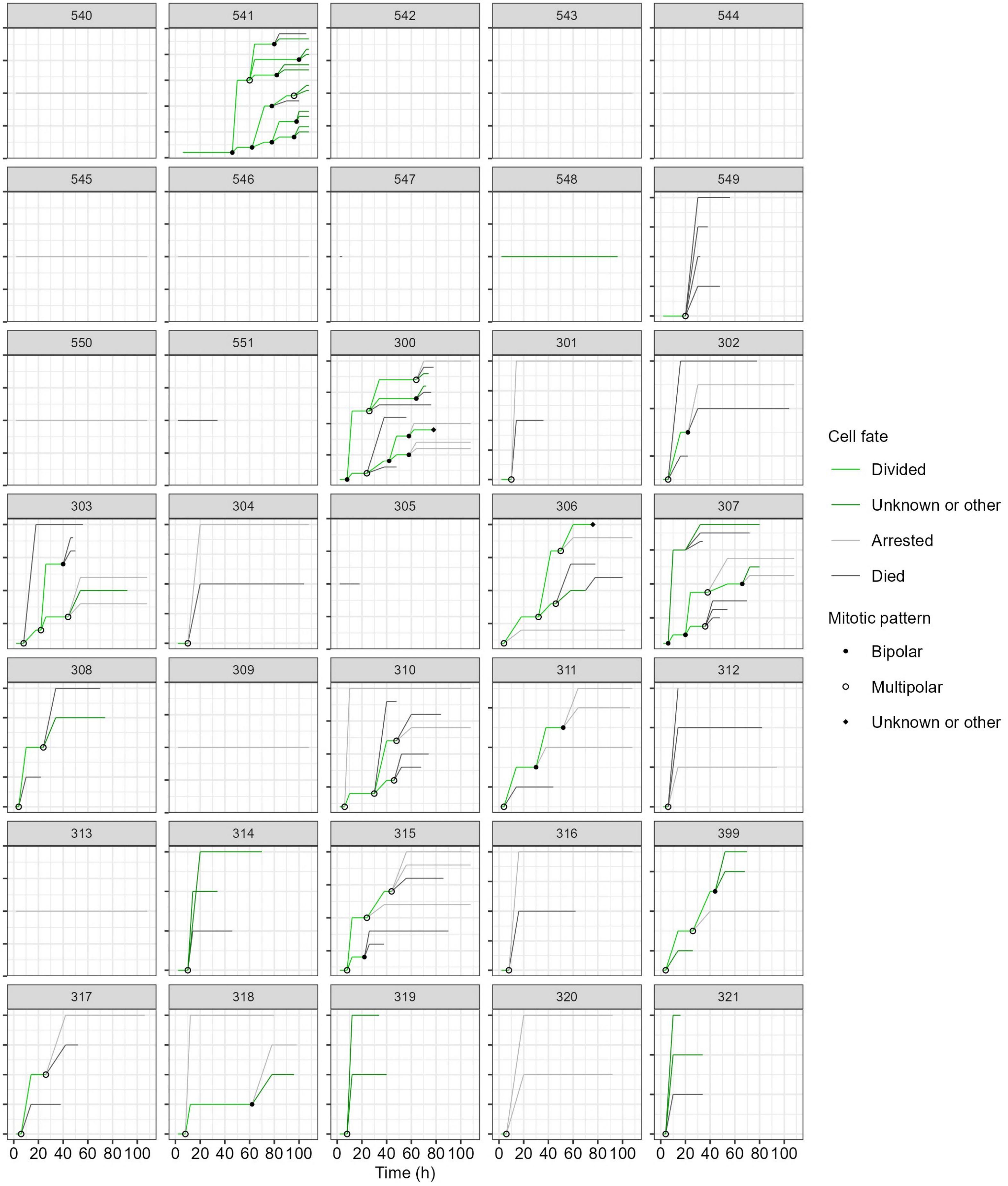

Supplementary material S1 (continued)

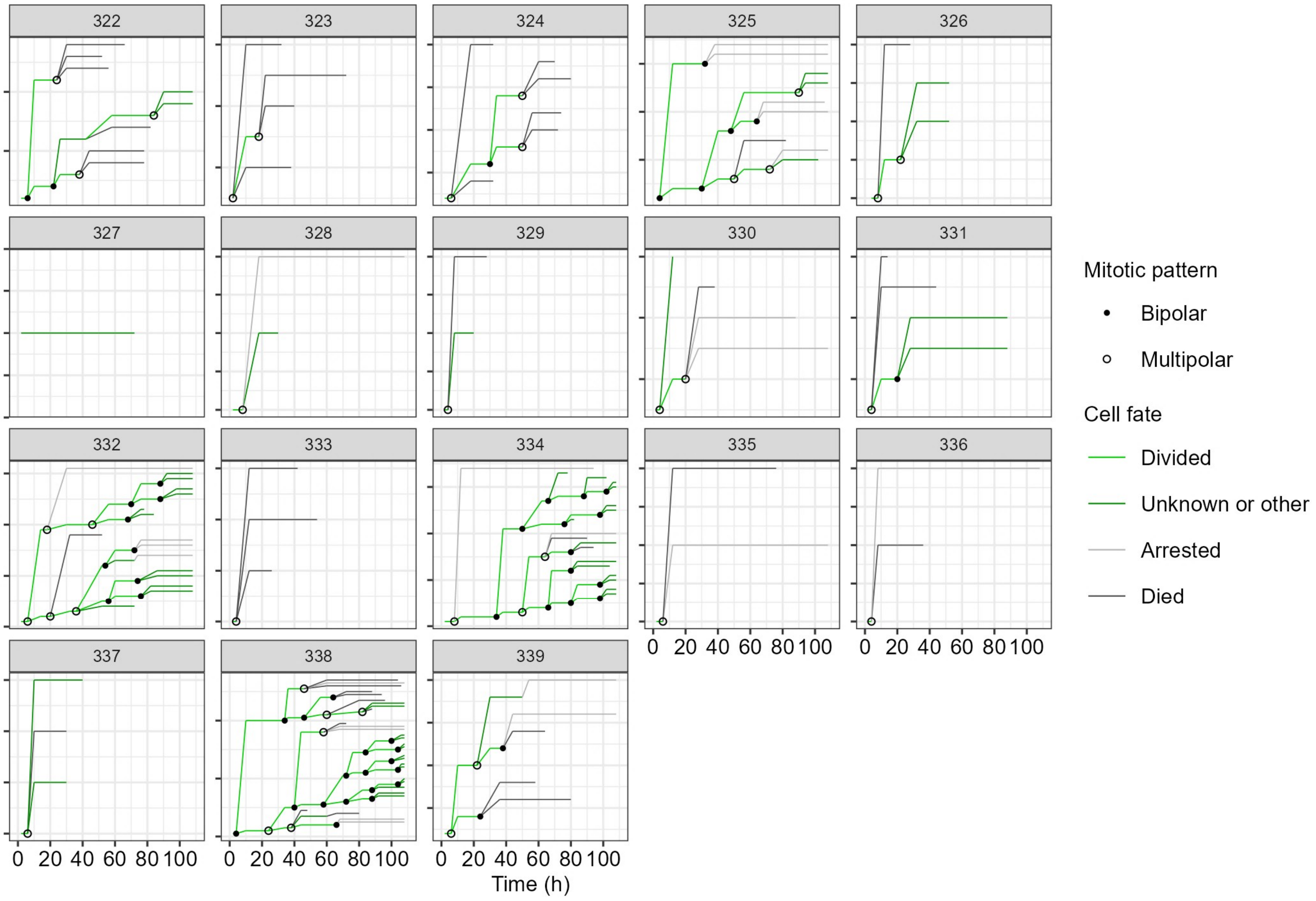
